# Directional selection rather than functional constraints can shape the G matrix in rapidly adapting asexuals

**DOI:** 10.1101/351171

**Authors:** Kevin Gomez, Jason Bertram, Joanna Masel

## Abstract

Genetic covariances represent a combination of pleiotropy and linkage disequilibrium, shaped by the population’s history. Observed genetic covariance is most often interpreted in pleiotropic terms. In particular, functional constraints restricting which phenotypes are physically possible can lead to a stable **G** matrix with high genetic variance in fitness-associated traits and high pleiotropic negative covariance along the phenotypic curve of constraint. In contrast, population genetic models of relative fitness assume endless adaptation without constraint, through a series of selective sweeps that are well described by recent traveling wave models. We describe the implications of such population genetic models for the **G** matrix when pleiotropy is excluded by design, such that all covariance comes from linkage disequilibrium. The **G** matrix is far less stable than has previously been found, fluctuating over the timescale of selective sweeps. However, its orientation is relatively stable, corresponding to high genetic variance in fitness-associated traits and strong negative covariance - the same pattern often interpreted in terms of pleiotropic constraints but caused instead by linkage disequilibrium. We find that different mechanisms drive the instabilities along versus perpendicular to the fitness gradient. The origin of linkage disequilibrium is not drift, but small amounts of linkage disequilibrium are instead introduced by mutation and then amplified during competing selective sweeps. This illustrates the need to integrate a broader range of population genetic phenomena into quantitative genetics.

## Introduction

Natural selection acts on multiple traits simultaneously. The mean trait value in a population can change either because of direct selection on trait X, or because of selection on trait Y plus a genetic correlation between X and Y (Lande 1979; Lande and Arnold 1983). These genetic correlations are described by the **G** matrix, which specifies both additive genetic variances and covariances. For example, in the case of two traits, each determined by an additive genetic component and an environmental component (*X* = *A*_*X*_ + *E*_*X*_ and *Y* = *A*_*Y*_ + *E*_*Y*_), then **G** is given by

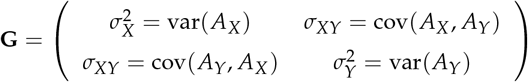

When the **G** matrix is stable, it is possible to infer past selection gradients (Lande 1979) and forecast future trait evolution (both direction and rate; Via and Lande 1985; Arnold 1992; Björklund 1996; Schluter 1996; Teplitsky *et al.* 2011, 2014). However, measurement of the **G** matrix has not come into widespread use for this purpose, perhaps in part because its stability cannot be assumed a priori. Theoretical models (Turelli 1988; Burger and Lande 1994; Jones *et al.* 2003, 2004), comparative studies (Björklund *et al.* 2013; Waldmann and Andersson 2000), and experimental evolution (Wilkinson *et al.* 1990; Shaw *et al.* 1995; Phillips *et al.* 2001) have all demonstrated that rapid change in the **G** matrix is possible. How stable **G** is in natural populations remains an open question (Steppan *et al.* 2002; Arnold *et al.* 2008).

Comparative studies suggest that at least certain aspects of **G** might be stable (Arnold *et al.* 2008). Early statistical approaches that compared the magnitudes of the individual variance and covariance elements suffered from multiple comparisons and lacked power (Shaw and Billington 1991; Brodie 1993; Carr and Fenster 1994; Roff and Mousseau 1999; Shaw and Billington 1991). Moreover, even statistically significant differences needed to be interpreted in the context of **G**’s geometric structure. More recent methods examine common principal components of **G** matrices and test for similarity in geometric shape, size and orientation, providing more biologically interpretable information (Arnold and Phillips 1999; Phillips and Arnold 1999). These methods suggest that the orientations of **G** matrices are often preserved between closely related populations (Arnold *et al.* 2008).

Our focus here is on traits highly correlated to fitness. Many of these are life history traits, which can also be highly correlated with one another. Specifically, they are often subject to constraints, e.g. acquisition vs. allocation, or competing allocations. The Charnov-Charlesworth model (Figure 1) describes how functional constraints among life history traits shapes genetic covariances (Charnov 1989; Charlesworth 1990; Walsh 2018, Chapter 35, Page 23). Functional constraints, where it is physically impossible for a phenotype to be simultaneously good in all trait dimensions (or more loosely, constraint-breaking mutations are vanishingly rare), can be visualized as a surface in trait space. Selection will quickly bring a population to this surface, but it may spread out along it when different points have similar overall fitness. From the population’s position along the surface, mutations can either decrease fitness in all dimensions (and be quickly lost), or they can increase fitness in some dimensions while decreasing it others (and potentially be retained as nearly neutral); mutation from the front is thus interpreted as subject to a pleiotropic trade-off. In this model, the cause of negative covariance among adaptive life history traits is constraint, and the stability of the constraint surface along which the spread occurs is thus thought to be the cause of the stable orientation of the **G** matrix.

**Figure 1.**
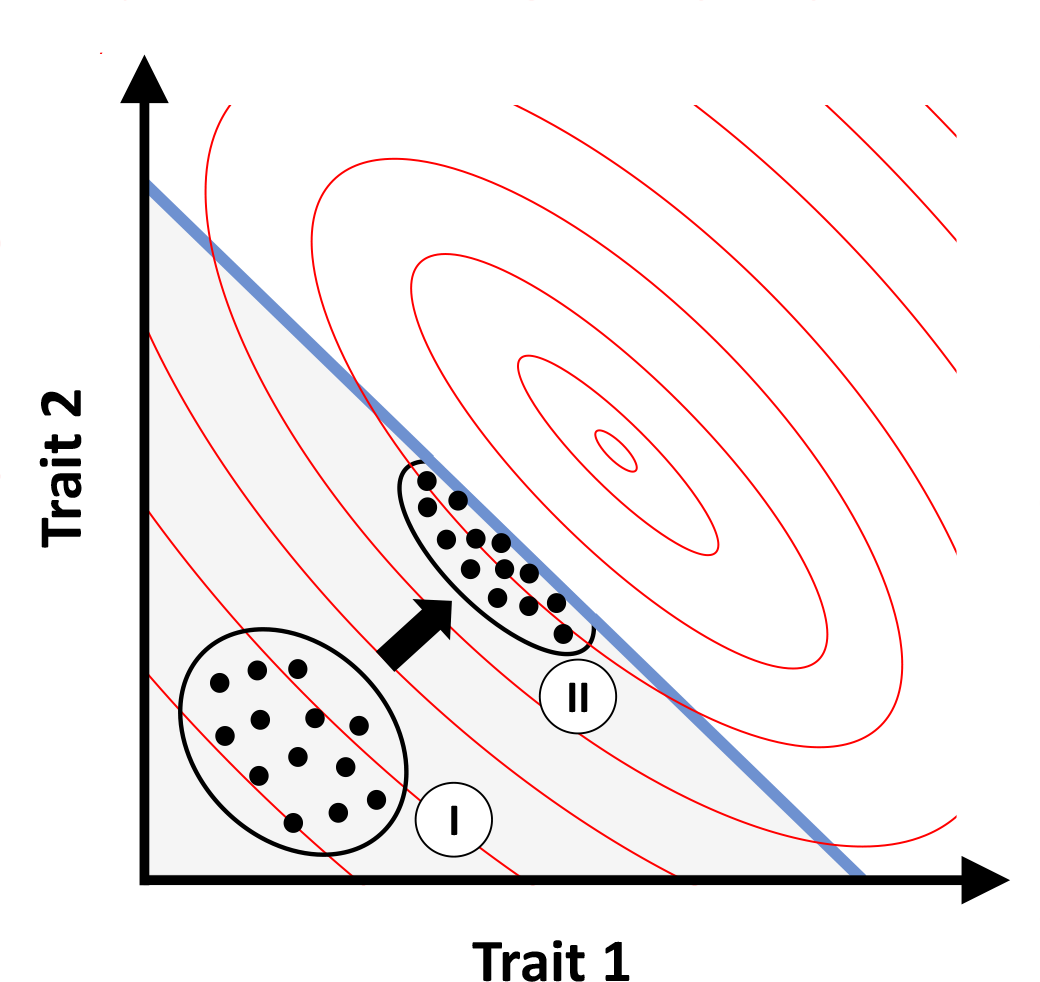
The Charnov-Charlesworth model, illustrated for selection on two traits subject to functional constraint. Functional constraint prohibits phenotypes above and to the right of the blue line. Fitness contours are shown as red ellipses, with highest fitness at the center. The population is initialized with trait values far from constraint (I). Selection then moves the population closer to the optimum, but functional constraint prevents the population from evolving past the blue line and it settles in mutation-selection balance at (II). Mutations with negative pleiotropy can be close to neutral and maintain trait variation along the line of functional constraint, while mutations affecting only one trait tend to reduce fitness and are purged.

In the alternative scenario that we consider here, there are no functional constraints on what trait values are possible. Indeed, we model the case of no pleiotropy, where each mutation affects only one fitness-associated trait. Every mutation creates a small amount of linkage disequilibrium; if the mutation is favored by selection, this linkage disequilibrium can be amplified by exponential growth of the mutant genotype, provided that there is insufficient recombination to interrupt this process. This amplified linkage disequilibrium then contributes to trait covariance if the alleles in linkage disequilibrium affect different traits. If there are multiple favored genotypes, each having an advantage in a different trait, then they may sweep simultaneously. We will show that during this process, there will be negative covariance between different fitness-associated traits. Instead of a physically impassable curve of phenotypic trade-off with negative covariance along it, we have a traveling wave with negative covariance (Figure 2). In this scenario, genetic correlations are due to linkage disequilibrium instead of to pleiotropy, the latter being absent from the model by construction.

**Figure 2.**
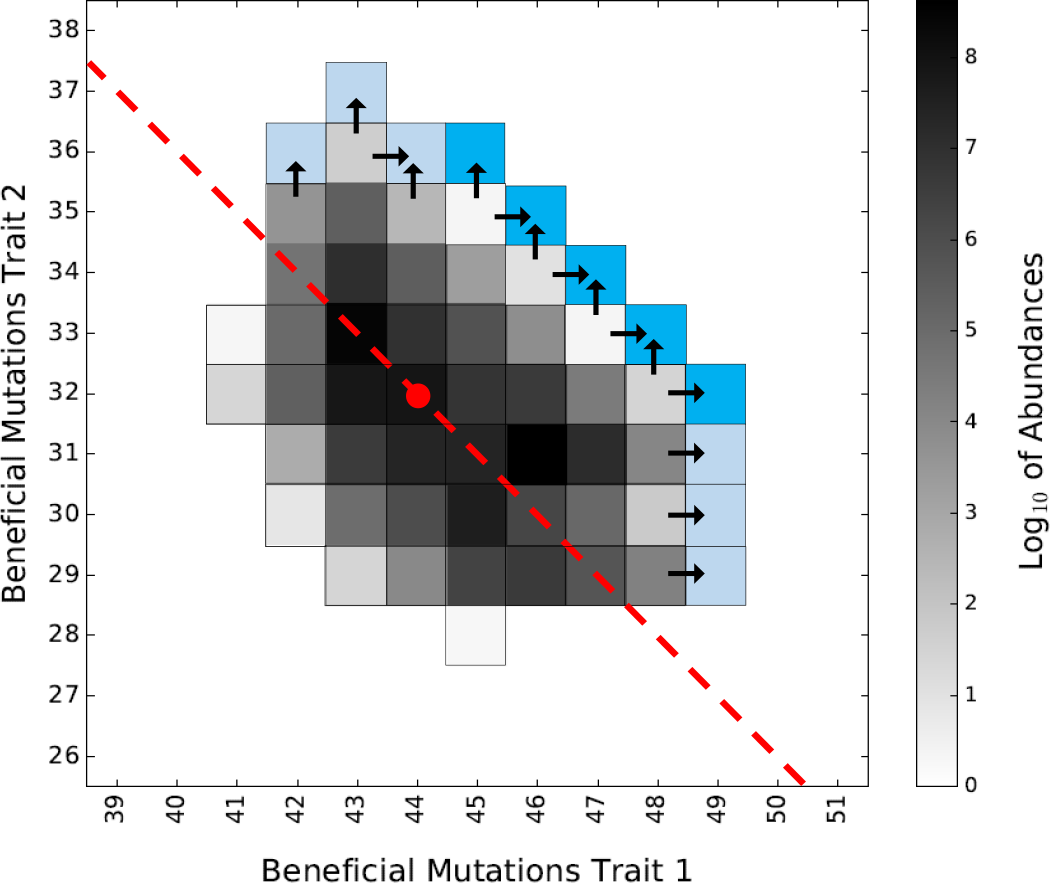
Representative two-dimensional genotype distribution. Individuals with equal numbers of beneficial mutations in each trait are combined into classes (squares). Abundances of the bulk (grayscale squares) behave deterministically, while abundance of the stochastic front (light and deep blue squares) behave stochastically. The fittest genotypes in the population (deep blue squares) are referred to as the high-fitness front. The red dot marks the class with approximately average numbers of beneficial mutations in each trait. Equally fit classes lie along fitness isoclines, which are the lines parallel to the red line shown. In asexuals, beneficial mutations must occur on the fittest genetic backgrounds in order to contribute to the adaptive process. These mutations are represented by arrows from the bulk into the stochastic front. Classes in the stochastic front become part of the bulk once their abundances have become sufficiently large, advancing the stochastic front to include new classes. As classes below the population mean fitness isocline decline and those above increase exponentially, a two-dimensional traveling wave is produced. Simulation parameters: *N* = 10^9^, *s* = 0.02, and *U* = 10^*-*5^.

Pleiotropy, rather than linkage disequilibrium, is often assumed to be the cause of most genetic correlations (Lande 1980a; Wagner 1989; Schluter 2000). Historically, a dominant role for pleiotropy was favored by the Edinburgh school of quantitative genetics, who argued that recombination would quickly eliminate linkage disequilibrium (Falconer 1993; Fox and Wolf 2006, Chapter 20), particularly in the randomly mating animal populations that the Edinburgh school focused on (Lande 1979, 1980b; Arnold *et al.* 2008; Robertson 1963). In contrast, quantitative geneticists in the alternative, Birmingham school largely worked on inbred lines of plants, where the effects of linkage disequilibrium were impossible to ignore (Robertson 1963; Mather and Jinks 2013, Page 25). As the Edinburgh view became more influential, models for the long-term evolution of the **G** matrix focused on genetic correlations that are derived from pleiotropy (Lande 1980a; Wagner 1989; Jones *et al.* 2003, 2004). We know far less about the dynamics of genetic correlations arising from linkage disequilibrium. Here we will show how completely different processes can give rise to similar patterns as the Charnov-Charlesworth model illustrated in Figure 1.

Here, we focus on asexual populations, because they are subject to the strongest linkage disequilibrium due to lack of recombination. Indeed, empirical studies of **G** in life history traits among the parthenogenetic soil-dwelling nematode *Acrobeloides nanus* demonstrate large instabilities in **G** (Doroszuk *et al.* 2008). More definitively implicating linkage disequilibrium, Pfrender and Lynch (2000) measured temporal instability in the **G** matrix for life history traits of *Daphnia pulex*, and found a buildup of covariance during asexual propagation that disappeared upon sex. Another reason to focus on asexuals is that relatively asexual microbes numerically dominate the biosphere, impacting all fields of biology (McFall-Ngai *et al.* 2013).

To study the effects of linkage disequilibrium on **G**, we have to consider explicit alleles or genotypes, not just quantitative traits. Beneficial mutations are discrete not infinitesimal, and appear on distinct genetic backgrounds. Selective sweeps compete with one another, causing beneficial mutations to be lost in a process known as “clonal interference”. Clonal interference slows adaptation (Hill and Robertson 1966) unless recombination brings beneficial mutations together in the same genotypes, reducing negative linkage disequilibrium between them (Fisher 1930; Muller 1932). Many classical models of clonal interference considered only two loci each with two alleles, or excluded drift (Felsenstein 1974). More recently, traveling wave models have been developed to describe clonal interference among large numbers of loci, arising at arbitrary mutation rates (Rouzine *et al.* 2003; Desai and Fisher 2007; Park *et al.* 2010; Good *et al.* 2012; Neher *et al.* 2010; Fisher 2013; Rouzine and Coffin 2010, 2007; Rouzine *et al.* 2008). These models show that clonal interference can have an enormous impact on adaptation rates.

To date, traveling wave models have treated only the evolution of “fitness”; we adapt them to also consider the evolution of individual fitness-associated traits and their correlations. We begin with Desai and Fisher’s (2007) framework of fixed population size *N* and beneficial mutations at rate *U*, each with the same selection coefficient *s*. This yields the rate of adaptation (Desai and Fisher 2007, Equation 41)

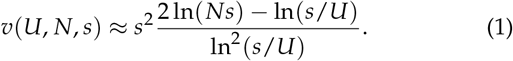

Equation (1) holds when *NU ≿* 1/ ln(*Ns*) and *U*/*s ≪* 1 (the “concurrent mutations regime” where beneficial mutations appear rapidly relative to the time required for any one of them to fix).

By Fisher’s fundamental theorem, *v*(*U*, *N*, *s*) approximately equals the additive genetic variance in fitness *σ*^2^; the approximation is due to the neglect of mutational flux (Desai and Fisher 2007). Thus, when there is only one adaptive trait, that trait’s additive genetic variance is given by *v*(*U*, *N*, *s*). With two adaptive traits, Equation (1) still gives the overall fitness variance *σ*^2^, but does not give its decomposition into the variances and covariance, i.e. it does not give the **G** matrix. Consider just two traits, each experiencing beneficial mutations at rate *U*. The overall adaptation rate is *v*(2*U*, *N*, *s*), and so by symmetry, adaptation in each trait alone occurs at rate *v*_1_ = *v*(2*U*, *N*, *s*)/2. This is lower than the rate *v*(*U*, *N*, *s*) that would occur if the other trait were not evolving. We do not know from this how the reduction in trait-specific adaptation rate is distributed in 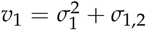 (see Appendix A for the derivation of this analog to the multivariate breeder’s equation). What is more, this distribution of adaptation rate reduction between 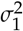 and *σ*_1,2_ might change over time i.e. **G** may not be constant.

Here we analyze a two-dimensional traveling wave model of asexual adaptation, and find that linkage disequilibrium alone, in the absence of pleiotropy, leads on average to greatly elevated variance of fitness-associated traits and to strong negative covariance between them. The **G** matrix arising in this way has a strong bias toward an orientation reflecting the direction of selection, rather than the nature of functional constraints, but its elements are highly unstable over the timescale of selective sweeps.

## Materials and Methods

We consider an asexual haploid population of fixed size *N* evolving in continuous time. There is no pleiotropy (each mutation affects only one trait), and there is no epistasis. To keep the model as simple as possible, we assume that there are two traits, each with the same rate of beneficial mutations *U*, and that each mutation has the same fitness benefit *s*.

We do not consider deleterious mutations (see Discussion). All individuals that have accumulated *i* mutations in the first trait and *j* in the second have the same Malthusian relative fitness *r*_*i*,*j*_ = *is* + *js*. We refer to a set of individuals with the same *i* and *j* as a “class” and denote their abundances and frequencies by *n*_*i*,*j*_ and *p*_*i*,*j*_ respectively. The population’s mean fitness is 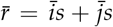, where 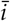 and 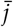 are the mean numbers of beneficial mutations (Figure 2, red dot). Equally fit classes lie on a fitness isoclines (Figure 2, red line). The selective advantage of an individual in class (*i*, *j*) with respect to an average individual in the population is 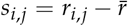.

We restrict our attention to the large *N* regime in which most classes are so large that their frequency grows or declines approximately deterministically due to selection (*N ≫* 1/*s*). We refer to these classes as the “bulk” (Figure 2, grayscale squares). However, higher fitness mutant lineages arising along the already high-fitness “front” of the population will start with a single individual, and initially behave stochastically (Figure 2, blue squares; arrows show mutations). Mutations creating new classes at the front will often go extinct before attaining appreciable frequencies, but some will make it to high enough abundances that they begin to grow deterministically. The transition from stochastic to deterministic behavior is called “establishment” and roughly corresponds to the lineage reaching abundance *n*_*i*,*j*_ *>* 1/*s*_*i*,*j*_ (Desai and Fisher 2007). Beneficial mutations that are not at the front are swamped by deterministic exponential dynamics and can be ignored.

Beneficial mutations at the front create new genetic variation, while selection in the bulk eliminates it. As the abundance of classes above and to the right of the red line grow in Figure 2, and those below and to the left decline, a two-dimensional traveling wave results, with the red line itself traveling diagonally up and right. The distribution of abundances within the bulk is the outcome of previous establishments at the front. Stochasticity in mutation and establishment at the front is thus propagated into the bulk (Hallatschek 2011; Fisher 2013; Desai *et al.* 2013; Pearce and Fisher 2018) and hence into the **G** matrix.

We simulate the dynamics of the two-dimensional traveling wave in discrete time using Matlab code developed by Pearce and Fisher (2018). They use a conservative threshold of *n*_*i*,*j*_ *>* 10/*s* for considering a class to be in the deterministic bulk for which stochastic fluctuations in growth can be ignored.

In the bulk, the growth (or decline) of a class due to selection is calculated according to

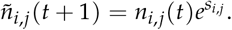

Following selection, abundances are adjusted for mutational flux, yielding

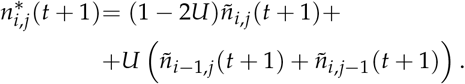

Outside of the bulk, to capture stochasticity due to drift and mutation, abundances are sampled from a Poisson distribution with mean 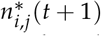. The classic Wright-Fisher model entails binomial sampling, but Poisson sampling is more convenient and a close approximation. Individuals with selective advantage *s* reproducing with birth rate 1 + *s* and death rate 1. Thus, each time step represents one generation.

To enforce a constant population size, abundances are rescaled by setting

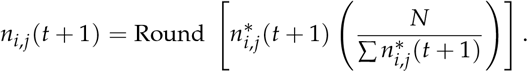

Simulations were initialized with monoclonal populations of size *N* and relative fitness set to zero. Parameter value ranges for *s* and *U* were informed by values obtained during experimental evolution: *U* ∼ 10^*-*5^ and *s* ∼ 0.02 for *Saccharomyces cerevisiae* (Desai *et al.* 2007) and *U* ∼ 10^*-*5^ and *s* ∼ 0.01 for *Escherichia coli* (Perfeito *et al.* 2007). Levy *et al.* (2015) used more sophisticated barcoding techniques to estimate the distribution of fitness effects for beneficial mutations in *S. cerevisiae*. Most lineages that established had selection coefficients in the range 0.02 − 0.05 with corresponding beneficial mutation rates on the order of ∼ 10^*-*5^. Although population sizes for *E. coli* during infection can be as large as ∼ 10^12^ (König *et al.* 1998; Wilson and Gaido 2004), we used smaller sizes of *N* ∼ 10^7^ *-* 10^11^ in our simulations. Before collecting data, we allowed the simulation to run for 5,000 generations to achieve beneficial mutation-selection balance.

We calculate trait means 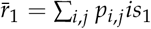 and 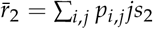, variances 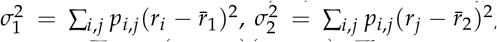, and covariance 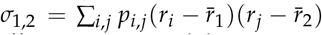. There are no environmental effects in our model. Because there is no epistasis, 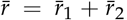, and the instantaneous rate of adaptation of the population is 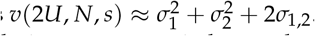.

Our simulations were carried on a desktop linux machine running Matlab 2018a. Our version of the Pearce and Fisher (2018) simulation code, together with Python scripts to generate figures, is available at https://github.com/MaselLab/Gomez_et_al_2018.

## Results

### 1. Clonal interference is substantial even for only two traits

As discussed in the Introduction, adding a new fitnessassociated trait reduces the rate of adaptation in a focal trait from *v*(*U*, *N*, *s*) to *v*_1_ = *v*(2*U*, *N*, *s*)/2. Generalizing to *k* traits, each with equal *U* and *s*, from Equation (1) we have

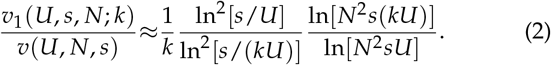

Each additional trait increases clonal interference on the focal trait by a diminishing amount, with curvature apparent even with respect to the logarithm of the number of traits (Figure 3). This suggests that much can be learned even from the simplest case of clonal interference between only two fitness-associated traits.

**Figure 3.**
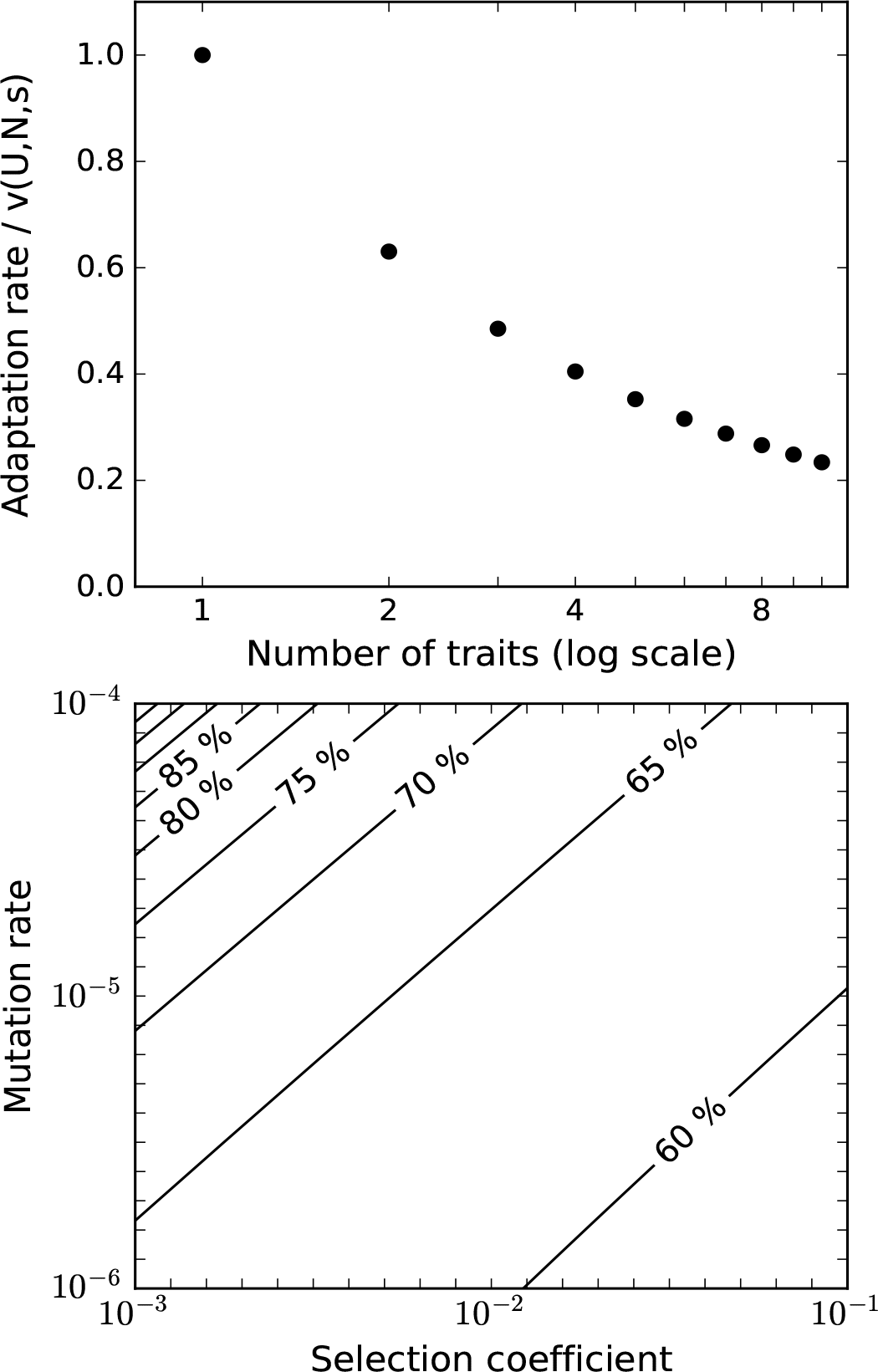
Clonal interference is substantial with even just a second adaptive trait. (a) The rate of adaptation of a focal trait as a function of the total number of traits undergoing adaptation, from Equation 2. The x-axis shows the total number of traits (including trait one) on a log scale. For *k* = 1, trait one evolves alone and there is no reduction, while for *k* = 2, trait one evolves at 62.2% of the rate that it would if it were not subject to clonal interference with a second trait. Steeply diminishing effects are seen with the addition of more traits. While the adaptation rate eventually asymptotes to zero for high *k* in Equation (2), note that this expression is only valid when *kU*/*s* ≪ 1. (b) Clonal interference between two traits is substantial and depends little on *U* and *s*, except in the top left corner where *kU*/*s* ≪ 1 has broken down. Contour lines are labeled as the rate of adaptation in a focal trait relative to the rate that would be achieved in the absence of clonal interference with a second trait, matching the second point in (a). *N* = 10^9^ throughout and the points in (a) used *s* = 0.02, *U* = 10^*-*5^.

### 2. The mean effect of clonal interference on G

The reduction in *v*_1_ due to clonal interference can be broken down into the effects on variance and on covariance, 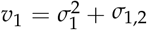, with clonal interference affecting the two components of **G**. We find that the reduction in *v*_1_ is driven by high levels of negative covariance (Figure 4, magenta circles). This negative covariance slows the removal of additive variance from the population, causing variance to be substantially higher (Figure 4, cyan circles). Negative covariance both cancels out the effect of the increased variance on the rate of trait change, and goes beyond it to cause the overall reduction in *v*_1_ to levels below *v*(*U*, *N*, *s*). While variances and covariance depend on *s* and *U*, the effects cancel out such that the reduction in *v*_1_ is insensitive to all three parameters.

**Figure 4.**
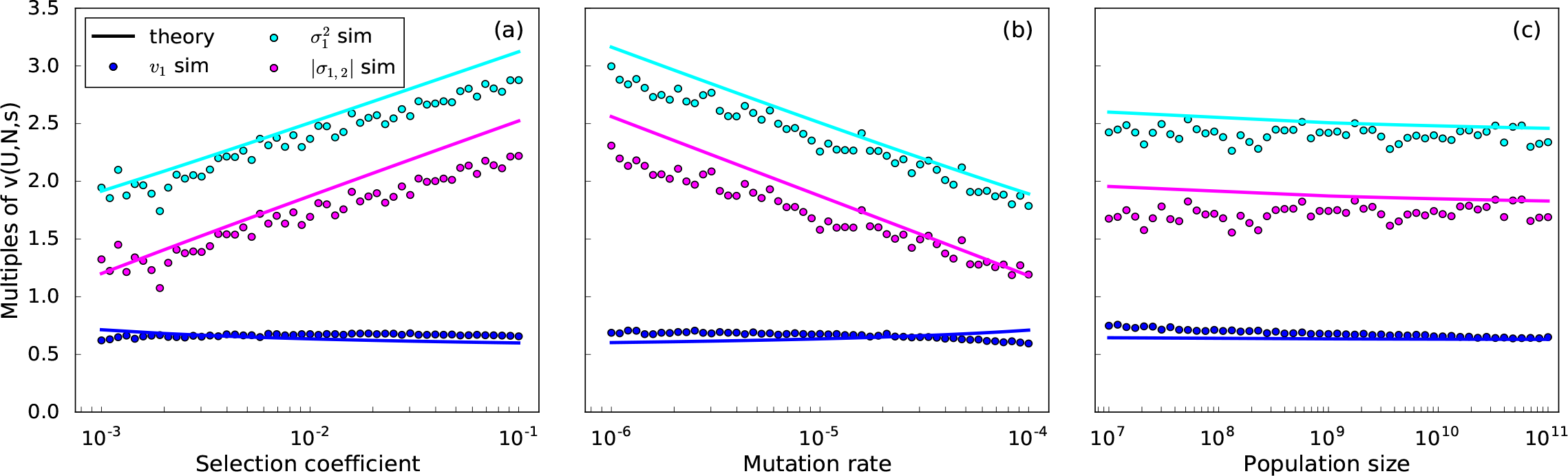
Variance 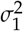 of a focal trait (cyan; simulated circles and Eq.(3)a solid line), the magnitude of its covariance *σ*_1,2_ with the other trait (magenta; simulated circles and Eq.(3)b solid line), and the trait’s contribution to adaptation *v*_1_ (blue; simulated circles and Eq. (1) solid line), averaged over 1.5 × 10^6^ generations. The y-axis is normalized relative to what the variance of the focal trait would be in the absence of the second trait; observed variance is always greater than this. While on its own, increased variance would accelerate adaptation, negative covariance more than cancels this out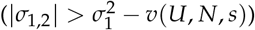 for a net reduction in the trait-specific adaptation rate below the value of *v*(*U*, *N*, *s*) that would be seen in the absence of clonal interference. For the parameter values not being varied on the x-axis, *s* = 0.02, *U* = 10^*-*5^ and *N* = 10^9^.

Appendix B uses the fitness-class coalescent developed by Walczak *et al.* (2012) to derive approximate analytic expressions for the expected values of 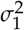 and *σ*_1,2_

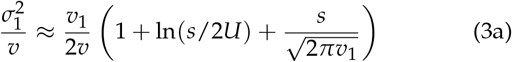

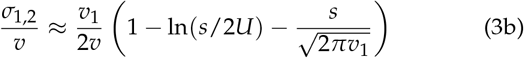

Equation 3 shows a good fit to simulations in Figure 4a-c.

Importantly, clonal interference in a rapidly adapting population can explain the common observation of high genetic variance in two fitness-associated traits combined with strong negative covariance between them, even in the complete absence of pleiotropic trade-offs imposed by functional constraints. This is striking, because under the Charnov-Charlesworth model, this is interpreted as evidence for constraint-driven trade-offs.

### 3. Variances and covariances are unstable

Figure 4 shows temporally-averaged variances and covariance. These values are highly unstable over time (Figure 5a). Indeed, the instability is so pronounced that variances and covariances in Figure 4 show significant noise, due to the difficulty in getting a good estimate of the mean even when averaging over a long time period. This is compatible with the fact that substantial instability in **G** has been observed empirically (Pfrender and Lynch 2000; Doroszuk *et al.* 2008).

**Figure 5.**
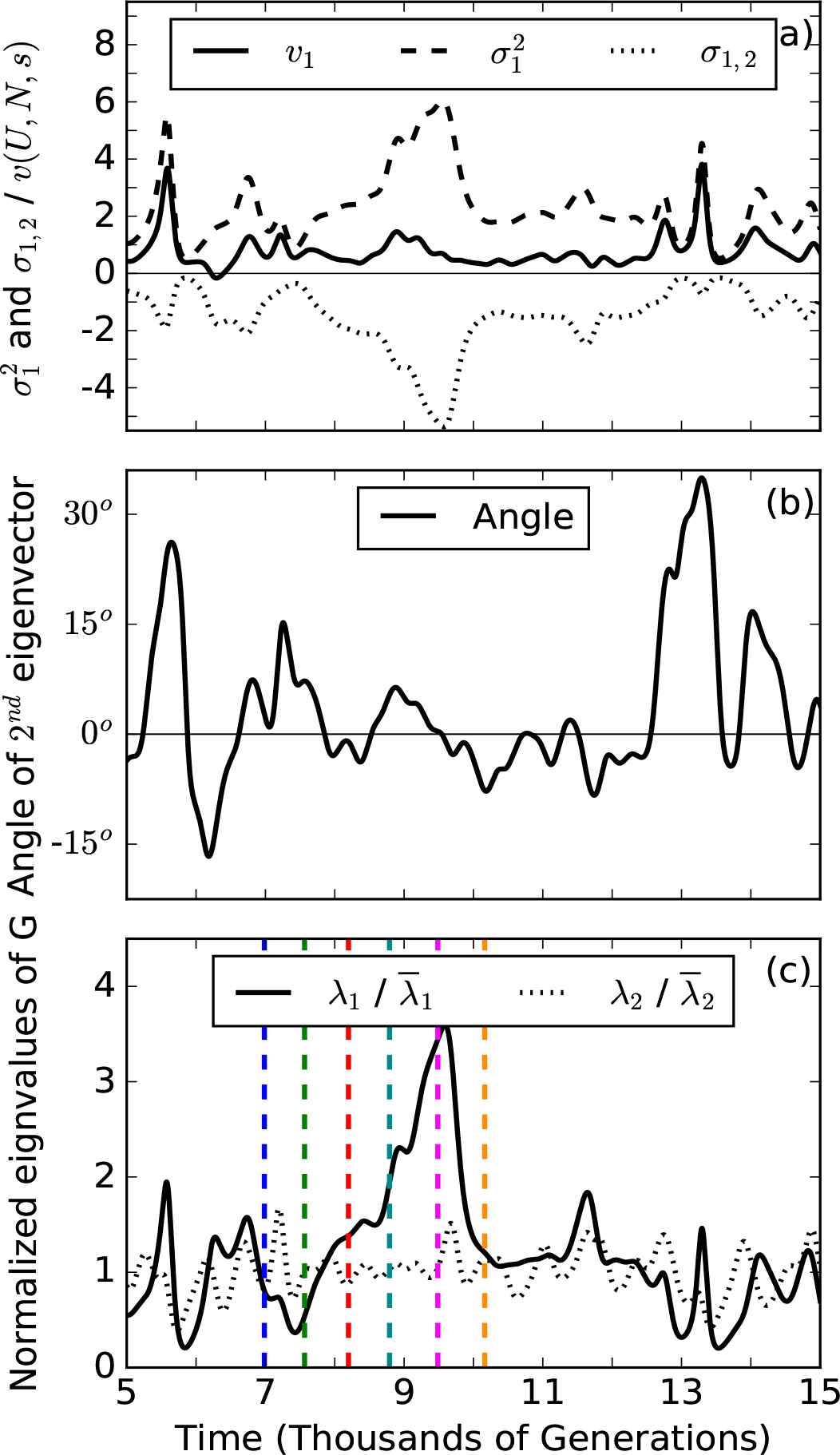
Behavior of **G** over time. (a) Variance of trait one (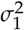, dashed line), its covariance with trait two (*σ*_1,2_, dotted line), and its contribution to adaptation (*v*_1_, solid line). Variances of 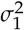 and *σ*_1,2_ (1.02) are about four times larger than the variance of *v*_1_ (0.27) (units of *v*(*U*, *N*, *s*)^2^). (b) The magnitude of the angle between the second eigenvector of **G** and direction of selection (vector (1, 1)) is on average 7.2° for the time period shown. Its grand mean is 0 and deviations from this mean orientation are usually small. (c) Normalized eigenvalues of **G**, *λ*_2_ and *λ*_1_, measure genetic variation along the direction of selection and the perpendicular “neutral direction”, respectively. Normalization disguises the fact that 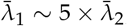; as a result, it is fluctuations in *λ*_1_ that drive those in variances, covariance, and angle. Fluctuations in *λ*_2_ have a different cause and are uncorrelated. Spikes in *λ*_1_ are due to enlargement of the high-fitness front, followed by its collapse (colored lines mark time points at which the 2D distribution is shown in Figure 7). Simulation parameters: *s* = 0.02, *U* = 10^*-*5^ and *N* = 10^9^.

Indeed, our simulations predict far greater instability than previously reported. Instability has been quantified in terms of change in the sum of the two variances. In previous simulations, this sum had a range of 80% of its mean over a period of 4000 generations (Jones *et al.* 2012). The same quantity in our simulations had a range of 192% of its mean over that same period of time. Our simulations also exhibited a much larger range for the inverse eccentricity of **G** (where eccentricity is defined by the ratio of **G**’s smallest and largest eigenvalue *λ*_2_/*λ*_1_), 320% of the mean inverse eccentricity in contrast to 125% for Jones *et al.* (2012). What is more, the simulations of Jones *et al.* (2012) were performed with *N* = 1024 while ours were performed with *N* = 10^9^, and the way in which genetic drift enters that model means that unlike in our model, increasing *N* would substantially reduce the instability.

Variances and covariances are dominated by the distribution of abundances among the most abundant or “dominant” genotypes. When we collapse the two-dimensional distribution into a one-dimensional traveling fitness wave, as shown in Figure 6a, the set of dominant genotypes are found primarily within the peak, which have approximately completed their exponential growth and are about to begin declining in frequency. This single one-dimensional “fitness class” consists of all the genotype classes that lie along a diagonal fitness isocline. The fluctuations of the **G** matrix can be understood by focusing on the distributions of frequencies within diagonal isoclines.

**Figure 6.**
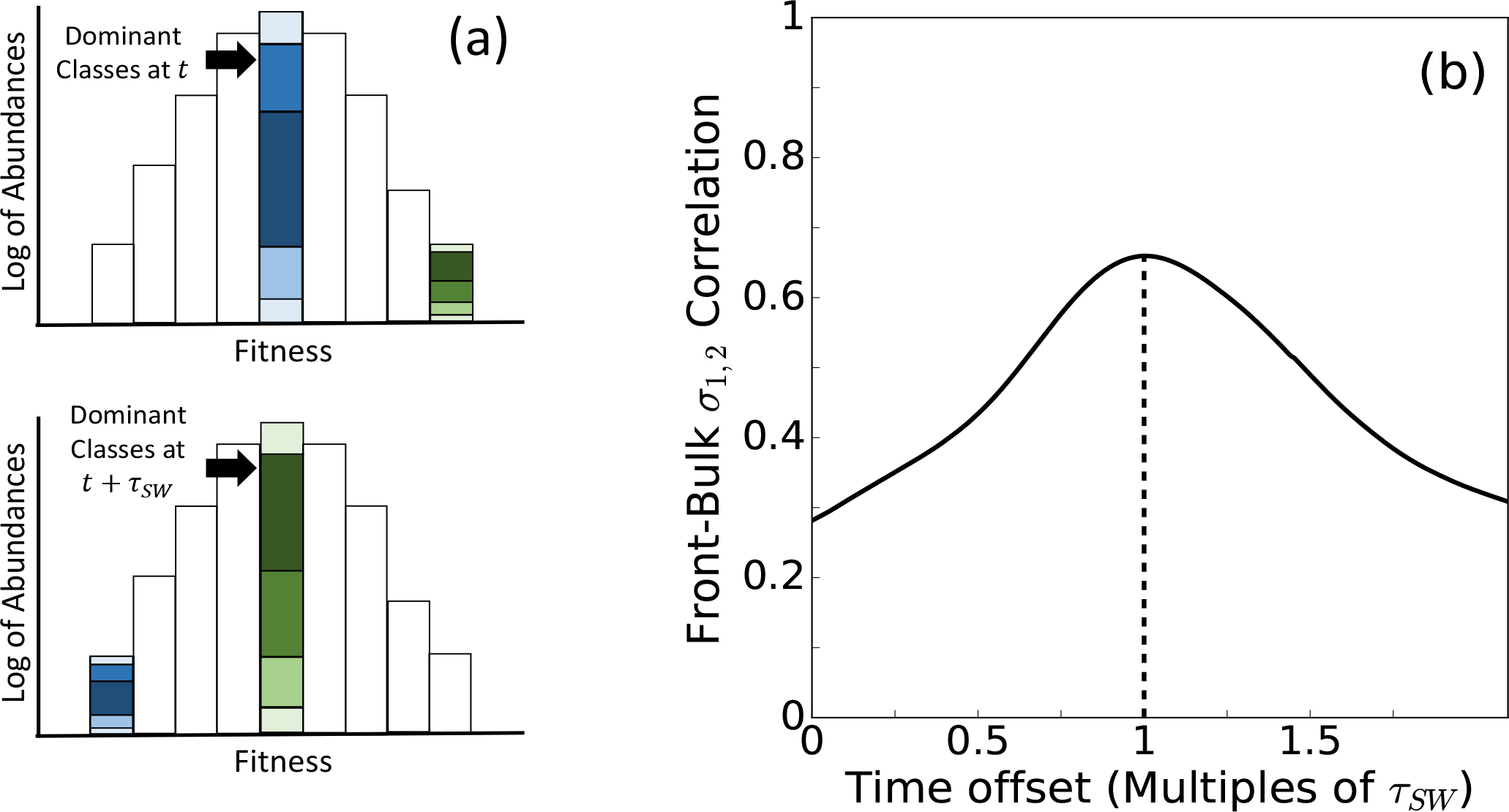
The **G** matrix is dominated by the highest abundance classes, whose composition reflects the previous distribution of classes along the high-fitness front. (a) Each fitness class combines all genotypes along the same fitness isocline (diagonal in Figure 2). The two-dimensional traveling wave in two-dimensional trait space can thus be projected onto a one-dimensional traveling wave in fitness space. Shading indicates the distinct genotypes defined in 2-dimensional trait space. Selection makes the most abundant fitness class exponentially larger than other fitness classes, meaning that the distribution of distinct genotypes within a single fitness class dominates the variances and covariances of the population as a whole. As the peak shifts from one fitness class to the next, variances and covariances may change substantially. (b) The correlation over time between covariance within the high-fitness front and covariance within the peak classes is highest with a time offset equal to the mean sweep time *τ*_*SW*_ given in Equation (4) (the average time required for the front to become the peak; dashed line). This is because the distribution of genotypes within a fitness class was set during the stochastic phase, and simply propagated deterministically until this fitness class became the most abundant. The dynamics of the stochastic front explain 66% of fluctuations in the covariance detected in the bulk after 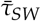 generations.

As discussed in the introduction, negative covariance in our model arises from the amplification of linkage disequilibrium generated by the beneficial mutations producing the fittest genotypes. Negative covariance thus originates with the stochastic dynamics of the high-fitness front. The relative ratios among high-fitness front classes are approximately “frozen” during the amplification that takes place after establishment, because beneficial mutations that occur after establishment of the high-fitness front (Figure 2, pale blue squares) contribute little to the relative frequencies of classes along a fitness isocline (Desai *et al.* 2013). As a result, the relative frequencies along a diagonal after establishment (Figure 6a, top green) are later found in the dominant classes (peak in Figure 6a, bottom green) once the traveling wave has moved that far.

The average time required for the high-fitness front to become the dominant group is given by the mean sweep time (Fisher 2013; Desai and Fisher 2007, Page 1178)

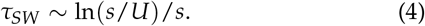

Figure 6b plots the correlation between covariances in the bulk and covariances in the high-fitness front as a function of the time offset between the two. The correlation peaks with approximately 66% of fluctuations in bulk covariance measured explained by the value of covariance at the high-fitness front *τ*_*SW*_ generations ago, confirming that fluctuations in **G** are primarily caused by changes in the distribution of relative frequencies of classes within successive high-fitness fronts. These have some short-term stability, because establishment times in the new front depend on the feeding classes that were part of the previous front. We shall see below that the instability of the components of **G** is driven primarily by fluctuations in the leading eigenvalue.

### 4. The orientation of the G matrix is mostly stable, while different forces drive the instability of the two eigenvalues

The eigenvalues and eigenvectors of **G** have been used to summarize its shape, size and orientation. Specifically, the orientation of **G** is specified by its eigenvectors, ranked by their eigenvalues, where each eigenvalue quantifies the genetic variance along its respective eigenvector. Each eigenvector can be specified by *m* angles relative to the *m* trait axes. Because the eigenvectors form an orthonormal set and thus each gives information about the others, only *m*(*m* – 1)/2 angles are needed for the matrix as a whole (Hohenlohe and Arnold 2008). Empirical comparisons of related populations often find that the orientation of **G** is stable even when its individual elements are not (Arnold *et al.* 2008).

For a two dimensional trait space, one can give the orientation of **G** using a single angle. Prior work on only two traits has used the angle between the first eigenvector and an arbitrary trait axis (Jones *et al.* 2003, 2004, 2007; Guillaume and Whitlock 2007; Revell 2007), or the angle between where the first eigenvector begins and where it is later (Björklund *et al.* 2013). We instead measure the orientation of **G** as the angle between **G**’s second eigenvector and the direction of selection (1, 1). By symmetry, the expectation of this angle is zero, with the expected eigenvectors of **G** being (1, –1) (first eigenvector) and (1, 1) (second eigenvector). The magnitude of the angle’s deviation from zero indicates the degree of instability in orientation, with 45° corresponding to a random matrix orientation. Since selection is identical on both traits, the vector (1, 1) in our two-dimensional trait space represents the direction of selection. To generalize our measure of orientation to more dimensions, we would measure the angle between the vector (1,…, 1) and whichever eigenvector is most closely aligned with this direction. We expect that this eigenvector will have the smallest eigenvalue since selection removes most genetic variation along in the direction of the vector (1,…, 1). We find that the angle measuring **G**’s orientation remains relatively stable. In Figure 5b the magnitude of this angle averages 7.2^*o,*^ which means that **G** remains closely aligned with the perpendicular “neutral” direction. This suggests that any observed stability in the orientation of **G** could reflect stability in the direction of selection of a traveling wave, rather than stability of functional constraints.

Figure 5c shows the behavior of the two eigenvalues, *λ*_1_ and *λ*_2_. The smaller eigenvalue *λ*_2_ measures genetic variation in the direction of selection, and *λ*_1_ measures genetic variation perpendicular to it, oriented along isoclines. Stochasticity in the speed at which the high-fitness front advances drives fluctuations in *λ*_2_, while fluctuations in the width of the high-fitness front drive fluctuations in *λ*_1_. In simulations, *λ*_1_’s average value over the period of the simulations was five times larger than the average *λ*_2_. It is the dynamics of *λ*_1_ that correspond to the fluctuations seen in 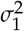 and *σ*_1,2_. In contrast, the dynamics of *λ*_2_ correspond to the overall adaptation rate *v*_1_ + *v*_2_, and to some extent also 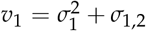 alone. This explains why, in our simulations, the variance in the time series of genetic variances and covariance is about four times larger than the variance in the time series data for *v*_1_ (Figure 5a).

As a new high-fitness front forms, it tends to be one class longer than the last front (Figure 2, dark blue classes), which will eventually increase *λ*_1_. When there is little variance in abundance among classes in the old front, beneficial mutations are fed into the new front at approximately the same rate, except for the two edge classes, which are fed at half the rate. Despite this disadvantage, classes at the edges do not on average take twice as long to establish, because the classes that feed them are growing exponentially. The probability that both edge classes establish before the next advance is therefore greater than the probability that neither will, creating an intrinsic tendency toward expansion of the high-fitness front (see Pearce and Fisher (2018) for a more detailed analysis of the front dynamics).

Over time, the abundances among classes in the front diverge stochastically. Small variations in abundance caused by stochastic establishment times change the rate at which beneficial mutations are fed into the next front, and thus cause establishment times to vary even more in the next front (Desai and Fisher 2007, Appendix D). Eventually, the differences in establishment times are large enough for the front to become segmented into competing sections that race to advance first. The winning section goes on to form a new and smaller front, as illustrated in Figure 7d to Figure 7e.

**Figure 7.**
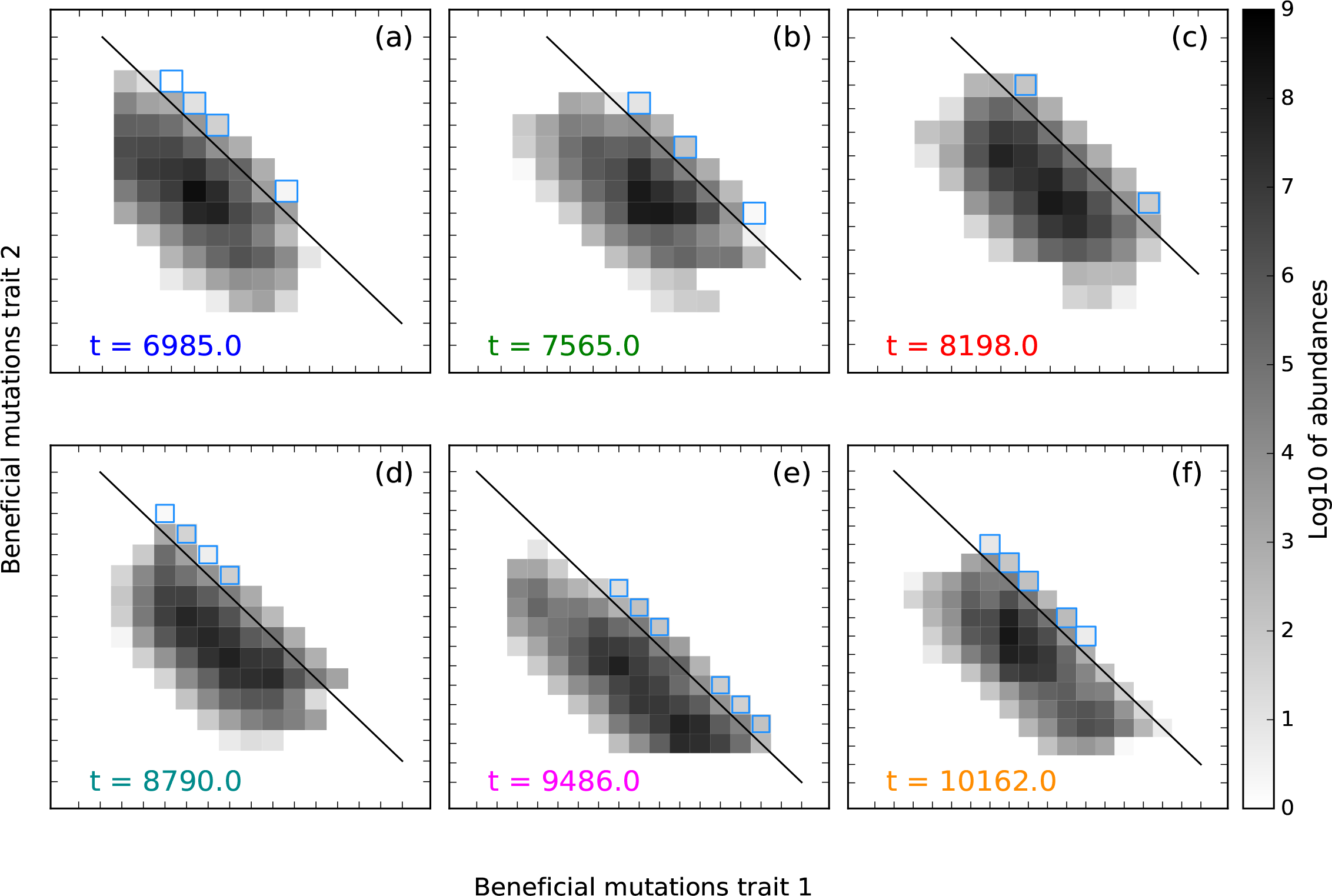
Expansion and collapse of the high-fitness front depicted with snapshots (a-f) of the two-dimensional distribution, corresponding by color to vertical lines in Figure 5b. Snapshots are approximately *τ*_*SW*_ apart (∼ 690 generations), in each case one generation before the first still higher fitness genotype appears by mutation. Squares with blue outlines are in the stochastic growth phase, and are fed mutations from classes along the fitness isocline indicated by the black line. (a) A narrow high-fitness front following a recent minor collapse, concentrated in two adjacent squares without outlines. (b) Even as genetic variation in the bulk declines, it increases within the narrow high-fitness front, breaking apart into two dominant segments. (c) The segments stochastically converge again, allowing for later widening of the front. (d) The width of the high-fitness front reaches a maximum, although maximum covariance will occur only *τ*_*SW*_ generations later. The lower right portion of the front is moving ahead of the top left section, setting it up for later collapse. (e) *λ*_1_ is at a local maximum, and the front is collapsing. The portion of the bulk no longer connected to the high-fitness front declines. (f) The high-fitness front collapses further as one segment continues to dominate its advances. Genetic variance in the neutral direction drops and causes negative covariance to decrease. Simulation parameters: *s* = 0.01, *U* = 10^*-*5^ and *N* = 10^9^.

Following collapse to a new, small front, variation in abundance among front classes is low, allowing for front expansion to resume until variation grows high enough to cause the front to collapse again. The high-fitness front cycles through phases of expansion and collapse (Figure 5c and Figure 7).

A different (and previously described; Desai and Fisher 2007) process drives fluctuations in *λ*_2_, namely instabilities in the rate at which the front advances, rather than instabilities in the width of the front. The value of *λ*_2_ is closely related to the distance *s*_*i*,*j*_ between the high-fitness front and the mean population fitness. Since the front advances stochastically, it will sometimes advance faster than the population mean fitness, temporarily increasing *s*_*i*,*j*_. This causes the front to accelerate, because fitness classes along the front will have a greater fitness advantage, and therefore produce more mutants with greater chance of establishment. Thus, *s*_*i*,*j*_ is dynamically unstable in the short-term, and so too is *λ*_2_. Eventually, fluctuations in *s*_*i*,*j*_ that accelerate the front also begin to accelerate the rate of adaptation in the bulk, once classes in the front become the dominant group *τ*_*SW*_ generations later. *s*_*i*,*j*_ then decreases, causing the front’s rate of advancement to decrease as well; this stabilizes *s*_*i*,*j*_ over the longer term.

## Discussion

Clonal interference in rapidly adapting asexual populations increases genetic variance in fitness-associated traits and creates strong negative genetic covariances between them, even in the total absence of pleiotropy. These effects are driven by linkage disequilibrium introduced by beneficial mutations at the high-fitness front and then amplified during selective sweeps. While the overall pattern is of high variance and strongly negative covariance, and the orientation of **G** is stably aligned with the direction of selection, the magnitudes of these **G** matrix elements are unstable over the timescale of selective sweeps. This pattern of wildly unstable magnitudes has been observed empirically in asexuals (Pfrender and Lynch 2000; Doroszuk *et al.* 2008); here we capture it for the first time in a formal model. This resulted in much larger instabilities in **G**’s components than had been found in previous models (Jones *et al.* 2012).

While this is a proof of principle work, we can nevertheless make a first attempt at qualitative predictions. The phenomenon described here predicts large fluctuations in the magnitude of **G** elements over a characteristic timescale of selective sweeps. In contrast, models in which adaptation has hit a wall of functional constraint predict small fluctuations only.

### The effects of deleterious mutations and recombination

In our model we made a number of simplifying assumptions. Two seem worthy of discussion: our neglect of deleterious mutations and of recombination. Deleterious mutations are undoubtedly common in all species, and even microbes can occasionally undergo recombination. However, we do not believe that a fuller treatment of either would lead to qualitatively different results from those presented here, which all rely on the same basic dynamic of the selective amplification of beneficial mutations at the high-fitness front. Deleterious mutations generally do not prevent new and fitter genotypes from appearing and sweeping. So long as these sweeps occur, the amplification of stochastically created linkage disequilibrium would continue to produce instabilities in **G**. The net behavior of **G** will be a combination of the dynamics we describe due to sweeps and the dynamics due to contributions from stabilizing selection at the trait level (as discussed in the Introduction). The contribution of this paper is to characterize the effect of sweeps on **G**, as a novel phenomenon.

Similarly, recombination will only eliminate the basic dynamics behind our findings if it is strong enough to eliminate linkage disequilibrium between genotypes generated by different beneficial mutations. Recombination, like mutation, can produce new fitness classes at the high-fitness stochastic front when. These too are amplified by subsequent selection, with qualitatively similar dynamics to the sweeps initiated by beneficial mutations as described by our model. With recombination, the genome will fragment into independently evolving linkage blocks, with an effective mutation rate *U*/(*RT*_*c*_) where recombination is represented as a larger total map length *R*, and *T*_*c*_ is the coalescence time (Neher *et al.* 2013; Good *et al.* 2014; Weissman and Hallatschek 2014). So long as *NU*/(*RT*_*c*_) ≫ 1/ ln(*Ns*), we expect partial selective sweeps and large negative correlations within linkage blocks, with negligible covariance across blocks. However, while the dynamics we describe will still occur, because genome-wide variance but not covariance scales with the number of linkage blocks, this will significant reduce the amount of covariance in **G**. While this reduces the scope of our findings, map length R may be negligibly small in the facultatively sexual world of microbes.

### Linkage disequilibrium in other models for G

Two previous quantitative genetic models have traced the impact of linkage disequilibrium on **G**. First is the Bulmer effect (Bulmer 1971; Walsh and Lynch 2018, Chapter 16). This describes the fact that selection which perturbs a previously stable state can generate linkage disequilibrium faster than it changes allele frequencies, at least when the number of loci contributing to a quantitative trait is large. This perturbation effect applies to much shorter timescales than the long-term steady state linkage disequilibrium considered here.

Second, Lande (1984) found conditions under which stabilizing selection on traits can create large and stable genetic correlations at equilibrium, via linkage disequilibrium. Selective sweeps do not routinely occur under stabilizing selection, but dominate in our regime. Because they amplify the stochasticity of mutation, they are responsible for the large instabilities in **G** seen in our work but not found in Lande’s. While many traits are undoubtedly both polygenic and under stabilizing selection (Charlesworth *et al.* 1982; Haller and Hendry 2014), it is also true that genomic evidence has made it clear that selective sweeps are abundant, even in sexual populations (Kern and Hahn 2018), and can make large contributions to linkage disequilibrium.

### Balance of forces versus mutation-driven evolution

Genetic variances and covariances are affected by selection, mutation, drift, recombination and migration (Walsh and Lynch 2018). These evolutionary processes have been historically viewed as “forces” acting on allele frequencies in the population (Gillespie 2010) and used to explain a variety of evolutionary phenomena. Past work on the **G** matrix has used the forces view both to gain intuition (Arnold *et al.* 2008), and to calculate equilibrium values of genetic variances and covariances (Lande 1975, 1980a, 1984; Tallis and Leppard 1988; Charnov 1989; Charlesworth 1990; Houle 1992).

The metaphor of “forces” evokes vectors that can be added. Equilibria of allele frequencies and related properties correspond to vectors that sum to zero. The classic example is a deleterious allele in balance between the force of mutation and the force of selection. The forces view has been criticized as a poor description of random genetic drift, which describes Brownian motion rather than a vector (Walsh 2000; Matthen and Ariew 2002; Walsh 2004), but it is still possible to talk about expected allele frequencies in balance between e.g. mutation, selection, and drift. A second, more serious critique is that mutation is not only a (very weak) vector acting on existing allele frequencies, it also controls which alleles exist to have their frequencies acted upon (Yampolsky and Stoltzfus 2001; Stoltzfus 2006).

In our context, interactions between evolutionary processes exhibit net effects that cannot be decomposed into independent vectors. This provides a third reason why the metaphor of evolutionary processes as vectors/forces is inappropriate. In our model, the most important stochastic “force” is not random genetic drift, but the introduction of beneficial mutations (combined with their establishment in the presence of drift). Each beneficial mutation appears on one genetic background, in complete linkage disequilibrium with all other loci. The contribution of this linkage disequilibrium to genetic covariance is small, because the allele frequency of a mutant initially present in a single individual is small. However, selection can dramatically amplify the magnitude of this linkage disequilibrium. The amplification of stochastic effects at the high-fitness front cannot be understood by a model of adding vectors, but arises through non-linear amplification. Fisher (2011) uses the metaphor of the mutational nose leading the dog to describe this scenario, whose effects on linkage disequilibrium are described by Garcia *et al.* (2018). No matter how big the total population is, the fittest subgroup steering the evolution of the population will always be small and hence stochastic, and its stochastic fate will eventually shape the entire population.

### Quantitative genetics versus population genetics

Our model differs in several important ways from previous quantitative genetic approaches. First, we assume no functional constraint nor other form of pleiotropy. Second, we assume no recombination, in contrast to quantitative genetic models that normally assume so much recombination such that linkage disequilibrium is negligible.

Third, most previous models used to simulate the stability of **G** have considered traits under stabilizing selection (Wagner 1989; Jones *et al.* 2003; Guillaume and Whitlock 2007; Revell 2007). A partial exception is Jones *et al.* (2004, 2012), who simulated a shifting optimum phenotype. While the moving optimum mathematically enters the model in the same way as stabilizing selection, it can mimic directional selection when the optimum is far from the population. However, their simulations focused on the regime where the population included the optimal phenotype. This ruled out the possibility of a stochastic nose and the amplification dynamics that we describe here. While the situation they considered was thus very different, they also found that directional selection increased genetic variances, and was responsible for stability in **G**’s orientation. However, their genetic covariance was positive, rather than negative, as a consequence of the shape of the phenotype-fitness map in their model.

The selective sweeps that are key to the phenomena we describe here are generally absent from most quantitative genetic models. Models for adaptation range on a spectrum from population genetics to quantitative genetics. Population genetic models include more details about allele and genotype frequencies and their temporal behavior. Quantitative genetic models are derived from population genetics, omitting details of genetic architecture in order to focus on statistical properties. More recently, population genetic details (in particular, a more discrete i.e. non-infinitesimal view of genes) have been essential to making progress in the historically quantitative genetic domain of QTLs and GWAS (Caballero *et al.* 2015; Simons *et al.* 2018). A deeper synthesis of the two approaches is not yet achieved.

Our model incorporates features from both methodologies. We specify discrete genotypes and track changes in their abundances, allowing us to model the establishment process and subsequent selective amplification of the linkage disequilibrium it produces, which translates into trait covariance under clonal interference. We used this to derive traditional quantitative genetic properties: traits, their genetic variances, and covariance. We found a classic pattern of high genetic variances and negative covariance in fitness-associated traits, but in our case it was not due to functional constraints. We also found that **G** is unstable over the timescale of selection, meaning that estimates of the **G** matrix are not likely to yield quantitatively accurate predictions for the long-term evolution of fitness-associated traits in the regimes considered here. Our work demonstrates the need to view adaptive scenarios from both a population genetic and quantitative genetic perspective, to better understand adaptive processes in traits that lie in between Mendelian and quantitative extremes.

## Acknowledgments

We thank Patrick Phillips and Bruce Walsh for helpful discussions, and Benjamin Good for the mathematics of Appendix B. Funding was provided by the National Science Foundation (DEB-1348262) and the National Institutes of Health (T32 GM084905).

### Appendix A

In our model of equal selection on two traits, the rate of adaptation in trait 1 is the sum of its genetic variance and genetic covariance, i.e.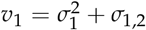. Here we derive this expression, which is analogous to the multivariate breeder’s equation but in continuous time and for a discrete genetic basis, from our definitions of mean fitness for each trait, 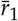 and 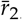, and the deterministic expressions for changes in genotypes abundances (see Methods). When the population is at capacity (∑_*i*,*j*_ *ni*,*j* (*t*) ≈ constant *N*), genotype frequencies within the bulk change according to the ODE

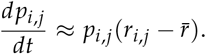

Substituting this into the time derivative of mean fitness in trait 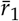 yields

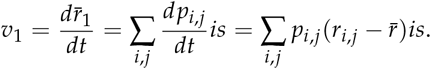

Substituting 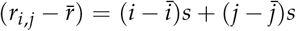 in the expression above gives

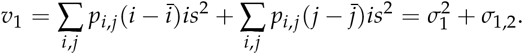

This can also be done for the second trait to get 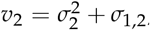, and thus, the total rate of adaptation must be

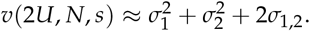

### Appendix B

The trait identity of each mutation (i.e., whether it influences trait 1 or 2) is a neutral marker in the present model. The differences between pairs of individuals in a single trait *k* = 1, 2 can be analyzed using a coalescent framework. Suppose that individual *i* has *n*_*i*_ mutations separating it from its common ancestor with individual *j*, and individual *j* has *n*_*j*_ mutations. Then we define indicator variables *I*_*k*,*i*_ (*l*) and *I*_*k*,*j*_ (*l*) that tell us whether mutation *l* occurred in trait *k* or not, so that the difference between the value of trait *k* in individual *i* vs. in the common ancestor is 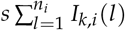. We then express the variance 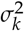 in one fitness-associated trait in terms of two randomly drawn individuals *i* and *j*:

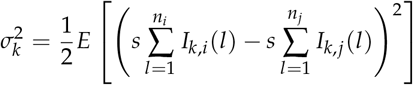

The factor of 1/2 comes from the fact that the expected squared difference of independent and identically distributed random variables *X* and *Y* is equal to twice the variance of either since *E*[(*X - Y*)^2^] = *Var*(*X*) + *Var*(*Y*) = 2*Var*(*X*). Rearranging,

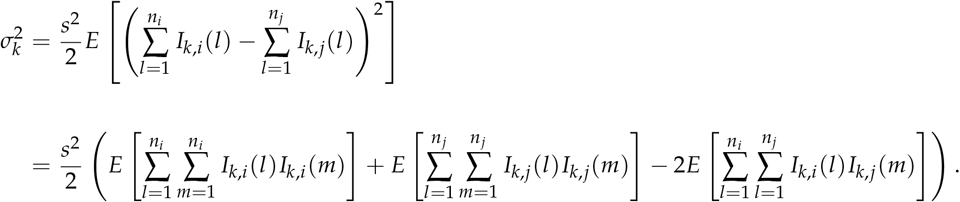

Calculating the expectations of these sums across binomial distributions of *n* trials, each with success probability 0.5, provides

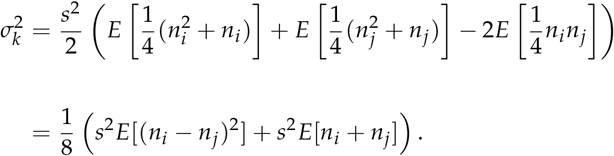

We note that the first term *s*^2^ *E*[(*n*_*i*_ − *n*_*j*_)^2^] = *E*[(*sn*_*i*_ − *sn*_*j*_)^2^] = *Var*(*sn*_*i*_) + *Var*(*sn*_*j*_) = 2*σ*^2^ ≈ 2*v*(2*U*, *N*, *s*), where *σ*^2^ is the total variance in fitness and hence yields Fisher’s Fundamental Theorem. The second term *E*[*n*_*i*_ + *n*_*j*_] is the average total pairwise heterozygosity at positively selected sites for two randomly selected individuals, Π. Substituting these into the expression for 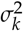 yields

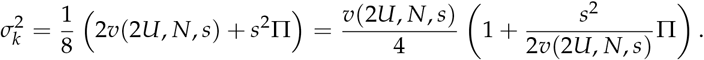

We obtain the covariance, *σ*_1,2_, using symmetry 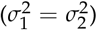 and solving for *σ*_1,2_ in the expression 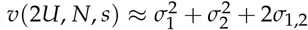 to obtain

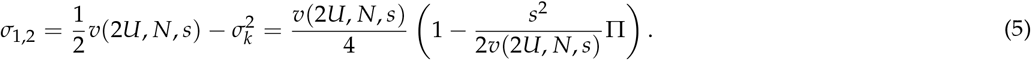

A expression for π is given by (Desai *et al.* 2013, Equation 30), derived from the fitness-class coalescent of Walczak *et al.* (2012). Expressed in terms of *v*(2*U*, *N*, *s*), we have

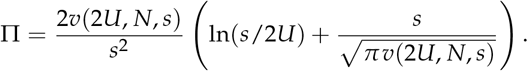

Substituting for π into the expressions for 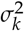 and *σ*_1,2_ provides

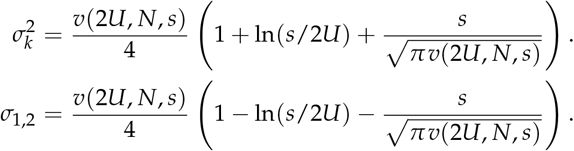

Finally, dividing both sides by *v* = *v*(*U*, *N*, *s*), and noting that 2*v*_1_ = *v*(2*U*, *N*, *s*), yields Equation 3. If we introduce the concept of additive genic variance *V*_*A*_ (additive genetic variance in fitness in the absence of linkage disequilibrium (Walsh and Lynch 2018, Page 550), then we can express trait variance and covariance in an alternative form. The comparison between *σ*^2^ and *V*_*A*_ (i.e. fitness variance in the presence vs. absence of linkage disequilibrium) has been used by Good *et al.* (2014) and Sohail *et al.* (2017) to quantify linkage disequilibrium. It can be expressed in terms of pairs of fitness-associated alleles (Lynch and Walsh 1998, Page 102) as

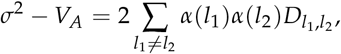

where *l*_1_ and *l*_2_ are segregating fitness-associated alleles, the *α*’s are the average effects of allelic substitution, and the 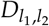 are the coefficients of linkage disequilibrium. The expected heterozygosity π can be written as

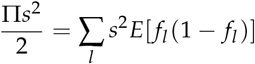

where the sum is over all segregrating sites *l* and *f*_*l*_ is the frequency at site *l*. The right hand side is the additive genic variance in fitness, *V*_*A*_ (Good *et al.* 2014; Walsh and Lynch 2018). We substitute both *v*(2*U*, *N*, *s*) = *σ*^2^ and 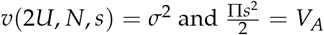 into Equation 5 to obtain

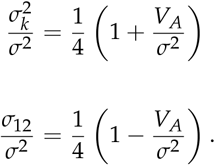

This describes how trait variances and covariances depend on linkage disequilibrium, which is quantified by the ratio *V*_*A*_ /*σ*^2^.

